# A range expanding ecosystem engineer influences historical and expanded habitats via the same causal pathways

**DOI:** 10.1101/2024.01.31.578099

**Authors:** Michael S. Roy, David Samuel Johnson, Jarrett E. K. Byrnes

## Abstract

Species are shifting their ranges in response to climate change. There remain many unknowns about relative impacts of range-expanding ecosystem engineers between historical and expanded habitats, however. The mud fiddler crab *Minuca pugnax* (=*Uca pugnax*) is shifting its range northward likely due to increased warming in the Gulf of Maine. A burrowing crab, *M. pugnax* affects ecosystem functioning in salt marshes south of Cape Cod, Massachusetts with unknown effects in expanded marsh habitats over 150km to the north. We therefore studied the *M. pugnax* range expansion to determine the extent that range expanding ecosystem engineers are influencing ecosystem functioning expanded ranges *relative to* historical habitats. We installed in 2017 and 2018 a series of crab-inclusion cages at both the UMass Boston Nantucket Field Station (historical range) and the Plum Island Estuary Long Term Ecological Research site (PIE-LTER, expanded range). For each site, year, and block, we measured in the beginning and end of the three-month experiment metrics of sediment strength, primary production, and decomposition. We developed and tested causal models using structural equation modeling (SEM) to determine direct and indirect effects of fiddler crabs on ecosystem functions. Despite site, year, and block variability, local environment influenced burrow density, which directly affected sediment strength and indirectly affected primary production in both ranges. Overall, understanding range-expanding ecosystem engineers in historical ranges was predictive for how they influence expanded habitats, despite inter-site heterogeneity. Therefore, it is critical to study relative impacts of range-expanding ecosystem engineers to understand total impacts of global range shifts.

## INTRODUCTION

Climate change is drastically altering physical, biological, and ecological processes globally. Higher average air and sea temperatures due to human caused fossil fuel emissions can contribute to more frequent and severe storms, induce sea level rise, shift phenology, and change historical species distributions (Brooker et al. 2007, Pitt et al. 2010, Langer et al. 2013, IPCC 2014, Yamaguchi et al. 2022). We focus on species range shifts, which are modifying marine ecosystems globally, including disruptions in both biodiversity (Edgar et al. 2005, Sorte et al. 2010), and ecosystem functioning at local and global scales (Gedan et al. 2009, Sorte et al. 2010, Balvanera et al. 2014, Hu and Juan 2014). Range-expanding ecosystem engineers could have especially large impacts as they modify their surroundings across time and space (Jones et al. 1994, Hastings et al. 2007, Crooks 2009). Despite their potential to influence ecosystem processes, few studies have looked at the impact of such range-expanding ecosystem engineers on both their historical and expanded ranges simultaneously (but see Aguilera et al. 2020). Therefore, a critical challenge remains to understand how range-expanding engineers affect ecosystem processes in expanded ranges *relative to* historical habitats, both currently and in the future (Gervais et al. 2021). Here we explore the direct and indirect consequences of the range expansion of an iconic ecosystem engineer, the fiddler crab *Minuca pugnax* (=*Uca pugnax*, Smith), to determine if its effect on ecosystem functioning is consistent with its historical range or wholly unique.

*M. pugnax* was historically located from southern Cape Cod, MA (hereafter referred to as “the Cape”) to Florida. In 2003, however, individuals were found in Scituate, MA (Sanford et al. 2006). By 2014, *M. pugnax* was documented in the Plum Island Estuary (PIE) (Johnson 2014), and has continued moving north (Johnson, *unpublished data)*. Mean annual temperatures of the GoM are rapidly increasing; the rate of warming in this ecosystem is 99% faster than the global average (Pershing et al. 2015) and is the likely mechanism driving the mud fiddler crab range expansion (Sanford et al. 2006). As an allogenic ecosystem engineer (i.e., a species that manipulates or otherwise changes the physical aspects of their habitat, Jones et al. 1994, Crooks 2009), *M. pugnax* burrows on average 15-25 cm (and can be as deep as 50 cm) into the sediment at the *Spartina alterniflora* dominated lower elevation zone of the salt marsh (i.e., the “low marsh”, Bertness 1985).

Previous research indicates that burrowing by the mud fiddler crab positively affects salt marsh ecosystem functioning in its historical ranges south of the Cape (e.g., Narragansett RI, Nantucket MA). By increasing sediment oxygenation rates, *M. pugnax* can facilitate aboveground growth of *S. alterniflora* live shoots, which in turn traps and deposits sediment for marsh vertical accretion (Bertness and Miller 1984, Bertness 1985), but not always (Williams and Johnson 2021). Alternatively, Martínez-Soto and Johnson (*Accepted, in Press*), via a control-impact study, showed that fiddler crabs reduced *S. alterniflora* biomass in PIE. Although they used historical data collected in separate research at the PIE LTER, they did not conduct a before-after-control-impact (BACI) study (i.e., direct before measurements were absent, Smith 2012). They therefore were unable to derive causal interpretations of their results, making it difficult to tell if crabs in their study were truly behaving differently in PIE or due to confounders in their design (Martínez-Soto and Johnson Accepted, in Press, Dee et al. 2023). Additionally, PIE crabs are both larger than their southern counterparts (Johnson et al. 2019), they can burrow in harder sediment than populations of *M. pugnax* in Nantucket (even after controlling for size, Wong et al. 2021), and PIE marshes are relatively different than Nantucket marshes (Gedan et al. 2011, Johnson et al. 2016). Therefore, it is unclear how *M. pugnax* will influence ecosystem functioning in its expanded range (either positively or negatively).

Given the abundance of literature examining the impact of *invasive* ecosystem engineers on their novel environment (Sueiro et al. 2013, Ma et al. 2021), bioinvasions ecology could provide needed insight on ecosystem impacts of recent native range expansions analogous to *M. pugnax*. Native range-expanders and invasive species can be similar in their impact, particularly if the range expansion occurs at relatively short time scales (i.e., years not decades, Sorte et al. 2010, Svenning et al. 2014). Several key differences between invasions and range expansions complicate this comparison, however. Bioinvaders often come from habitats many thousands of kilometers distant and have mixed impacts to invaded habitats (frequently negative, but sometimes positive or neutral, Castro et al. 2017, Pyšek et al. 2020, Vivó-Pons et al. 2020). Range-expanders, on the other hand, are seeded from ecosystems relatively close to their expanded habitats (Zacherl et al. 2003, Sorte et al. 2010, Bates et al. 2014). As such, the typical barriers (e.g., allee effects) and benefits (e.g., the enemy release hypothesis, lack of coevolutionary history with naïve prey) that an invasive species frequently encounters in a non-native habitat might not apply to range-expanding species in their expanded range (Colautti et al. 2004, Case et al. 2005, Blakeslee et al. 2013, Wallingford et al. 2020). Therefore, bioinvasions may not provide sufficient baseline context, underscoring the need to study how native range-expanding ecosystem engineers effect historical versus expanded habitats.

Using the mud fiddler crab as our model species, our goal was to determine relative impacts of range-expanding ecosystem engineers on ecosystem functioning between expanded and historical ranges. Our study specifically determined whether *M. pugnax* affects ecosystem functioning differently in Nantucket, MA (i.e., historical range) versus Plum Island Estuary, MA (i.e., PIE expanded range) and whether any observed effects were density dependent. We conducted fiddler crab inclusion experiments in both Nantucket and PIE in summers 2017 and 2018 and analyzed our results using Structural Equation Modeling (SEM) to determine pathways between fiddler crabs and each of our metrics of ecosystem functioning: 1) sediment strength (i.e., stability); 2) primary production; and 3) decomposition. Given strong site level differences, we expected that fiddler crabs influence these ecosystem functions differently in each location. Our research demonstrates, however, that fiddler crabs influence two of the three of our metrics of ecosystem functioning via the same patterns and processes in both historical and expanded ranges.

## METHODS

### Site Characterization

Folger’s marsh at The University of Massachusetts Boston Nantucket Field Station (NAN, 41.29368, −70.04005) and the salt marshes of the Plum Island Estuary Long Term Ecological Research (PIE LTER) site (PIE, 42.74642, −70.83701) are approximately 174 kilometers apart (see Supplemental Figure SF1) and differ in species richness, thermal regimes, sediment strength, sediment composition, and tidal range (Gedan et al. 2011, Johnson et al. 2016). As a burrowing species of crab, the survival of *Minuca pugnax* could be influenced by sediment hardness especially. In NAN, sediment strength is on average ~10psi (68.95 kPa) due primarily to silty oceanic inputs of sediment (*pers. obs*, Langlois 1979). PIE in contrast has a mean sediment strength at ~25-50psi (172.4-344.7kPa) due to primarily peaty organic rich sediment from low turnover of decomposing material and riverine inputs of sediment (Deegan et al. 2012, Vincent et al. 2013, Johnson et al. 2016, for a more detailed description of site differences between NAN and PIE, see Supplemental Methods 1). Therefore, such site differences made NAN and PIE ideal ecosystems to compare relative impacts of a range expanding ecosystem engineer on ecosystem functioning in historical versus expanded habitats.

### Caging Experiment

We conducted crab inclusion experiments from July to September/October in both 2017 and 2018, for both NAN and PIE. We manipulated fiddler crab densities in cages to create a blocked replicated regression design (Cottingham et al. 2005) with densities of 0-20 crabs at 4 crab intervals with random treatment placing, with 4 replications per treatment as well as a no cage control treatment (7 treatments x 4 replications = 28 plots per site per year). To quantify crab impacts, we measured a suite of plot properties including sediment strength (i.e., stability and compaction) at the beginning and end of the experiment, cordgrass (*Spartina alterniflora*) living shoot density at the beginning and end of the experiment, *S. alterniflora* biomass at the end of the experiment, fiddler crab burrow density at the end of the experiment, and rates of *S. alterniflora* belowground decomposition by the end of the experiment.

Prior to adding crabs to each fiddler crab inclusion cage, all crabs were removed from their burrows in each caging plot in a non-destructive manner. We established control plots per block to measure the same initial and final metrics of ecosystem functioning as caging plots but without a cage to determine degree of caging effects on our results. We compared these control plots to plots with ambient densities in NAN (~20 crabs plot^−1^, Aspey 1978) and in PIE (~4 crabs plot^−1^, Martínez-Soto and Johnson 2020) using ANOVA. Across all the metrics we measured, there were no significant differences between control plots and 20 crab cages in Nantucket (NAN), or between control plots and 4 crab cages in Plum Island (PIE), indicating the absence of a caging effect in this study (for a table of test statistics, see Supplemental Table ST1).

Cages were 0.25m^2^ x 1m and were constructed of 0.635cm polypropylene plastic mesh, ½ inch PVC pipes, and zip ties. In addition, each cage was buried ~12-15cm below the marsh surface to help prevent crabs burrowing out the bottom. To help prevent fiddler crabs from entering or exiting the top of each cage, we placed ‘flashing’ at the top ~8cm of the inner and outer sides of each cage. For 2017, we used heavy-duty packing tape as flashing due to cost; however, heavy-duty tape was missing by late summer in PIE (likely due to the meso-tides characteristic of this habitat). Therefore, when constructing the same experiment in 2018, we used steel plating, again at the top ~8cm of the inner and outer sides of each cage, which lasted the duration of the 2018 experiment. In all our statistical analyses, we controlled for year to account for this between year variability that may be caused by our 2017 flashing setup, or by interannual variability in environmental conditions common in marsh ecosystems. Our analyses indicate that while we observed significant differences between years, we largely observed the same patterns and processes driving our system for each year. Therefore, the failed flashing in 2017 does not appear to have had a qualitative effect on the overall results from this study (see Supplemental Tables for test statistics controlling for year). Regardless, we included year as a factor in all models to control for any possibility of multiple versions of treatments (Kimmel et al. 2021). We also carefully pruned or re-seeded crabs in cages by the middle of the experiment (~July/August for both sites and years) to maintain our manipulated fiddler crab densities per cage.

Prior to the addition of crabs, we measured our starting plot properties including initial *Spartina alterniflora* live shoot density in the top right corner of each plot (total area = 62.5 cm^2^) and initial sediment strength using a Humboldt Proctor style Penetrometer (model number H-4139), averaging sediment strength measurements among three corners of each plot (for 2018, sediment strength in 2017 was measured once in the center of the plot, see Supplemental Figure SF2 showing no difference between methods). We installed decomposition assays buried at 5cm and 15cm below the surface for each plot before adding crabs. In 2017, we used dried tea bags; however, all tea bags were completely disintegrated for each plot by the end of the experiment. Therefore in 2018, we buried 10.0g of air dried (20C for 24hours) live aboveground *S. alterniflora* material (collected from nearby creeks in each location) in mesh litter bags at our two decomposition depths. At the end of the experiment, we measured or remeasured: *S. alterniflora* live shoot density, final sediment strength, final burrow density, and aboveground biomass of *S. alterniflora*. We used *S. alterniflora* live shoot density and dry aboveground biomass collected at the end of the experiment (i.e., *S. alterniflora* biomass) as proxies for primary production (Angelini et al. 2015). We also excavated, rinsed, and dried the decomposition mesh bags in 2018, at 60°C to obtain a final litterbag mass of decomposed plant material. Lower final masses indicated more decomposed *S. alterniflora* from the baseline than higher masses (i.e., higher decomposition with lower masses).

### Statistical Analyses – Constructing and Evaluating our SEMs

To evaluate the direct and indirect effects of fiddler crabs, we developed three Structural Equation Models (SEMs, Bollen 1989, Grace 2006) for sediment strength, primary production, decomposition. We conducted all statistical analyses in R version 4.0.2 (R Core Team 2020) using the *piecewiseSEM* package (Lefcheck 2020) with generalized linear models (glms) as specified due to residuals from initial linear model fits with gaussian error violated assumptions of normality of residuals. All models were evaluated to meet assumptions of a uniform distribution of quantile residuals using the *DHARMa* package (Florian Hartig, 2020). Further, to cope with omitted variable bias and to make sure our models retained causal interpretation, we incorporated site, block, and year as fixed effects where specified (Bell et al. 2019). We assessed fit on all models using tests of conditional independence and Fisher’s C (Shipley 2000). We created three separate SEMs, rather than one grand model, to parse out model complexity and to better understand whether crabs interact with each ecosystem function individually. To access our data and scripts, please visit our github repository at https://github.com/MRoy120/Fiddler_SEM.

For our sediment strength Structural Equation Model (SEM), we built a model where initial sediment strength, crab treatment (i.e., initial crab density), and their interaction determined number of burrows. As burrows were an overdispersed count variable, we used a glm with a negative binomial distribution and log link function (O’Hara and Kotze 2010), evaluated for significance using Likelihood-Ratio (LR) chi-squared, Type II sums of squares. In our model, final sediment strength was then influenced by number of burrows, as a metric of crab activity, initial sediment strength, and the interaction between initial sediment strength and site. We fit this model with a Gaussian error and identity link, evaluated for significance using a Type II sums of squares F-test. As site, year, and block could possibly influence initial and final sediment strength as well as the ability of crabs to create burrows, we dealt with this omitted variable bias by incorporating year, site, and block into our models as a fixed effect (except in the case where site interacts with initial sediment strength, Bell et al. 2019).

For our primary production SEM, we used the sediment strength SEM described above as a baseline model. The primary production SEM model was built such that final live *S. alterniflora* live shoot density was determined by crab density, burrow density, initial sediment strength, and initial live shoot density. We also modeled whether initial live *S. alterniflora* live shoot density differed by sites, blocks, and years. As with burrows, both initial and final live shoot densities were overdispersed count variables; therefore we used a glm with a negative binomial distribution and a log link function for these two models as well (O’Hara and Kotze 2010). We evaluated both models for significance using Likelihood-Ratio (LR) chi-squared, Type II sums of squares.

In the case of *Spartina alterniflora* aboveground biomass, we built a model where crab density, burrow density, initial sediment strength, initial live shoot density, final live shoot density, and final sediment strength determined *S. alterniflora* aboveground biomass by the end of the experiment. In addition, we modeled the interaction between live shoot density at the beginning and end of the experiment (i.e., synergistic effects of live shoot growth from the beginning to the end of the experiment) affecting *S. alterniflora* biomass. Finally, we included the interaction between *S. alterniflora* final live shoot density and final sediment strength as a predictor of aboveground biomass to determine how bio-physical interactions impact *S. alterniflora* aboveground biomass. For *S. alterniflora* aboveground biomass, we fit a gamma distributed generalized linear model (glm) in the base R ‘*glm*’ function, again using the log link family function. As with the sediment strength SEM, we included site, block and year into our models as a fixed effect (Bell et al. 2019). We also evaluated this model for significance using Likelihood-Ratio (LR) chi-squared, Type II sums of squares.

As noted above, our decomposition assay failed in 2017, therefore only data for 2018 were used to construct our decomposition SEM. Due to limited power using only one year of data, we excluded connections from both our sediment strength and primary production causal models in our decomposition SEM. However, initial sediment strength and burrow density remained both as response variables and predictors in the decomposition SEM to determine whether sediment strength at the beginning of the experiment and burrows affected decomposition. In addition to burrows and initial sediment strength, depth (i.e., depth each litterbag was buried below the sediment surface), an interaction term between burrow density and depth, and an interaction term between burrow density and site were modeled where they affected decomposition. Decomposition (as measured by our final litterbag mass), was fit with a Gaussian error and identity link, and evaluated for significance using a Type II sums of squares F-test. All models passed statistical tests of assumptions except the model fit for *Spartina alterniflora* aboveground biomass, which included high variance inflation factors and a significant deviation from a Kolmogorov-Smirnov test. We corrected this model to eliminate these violations (see Supplemental Methods 2).

We tested each SEM for goodness of fit (i.e., p>0.05, no significant differences between observed and model generated variance-covariance matrices) and found that all three SEMs met the criteria for good model fitting (for a table of p values demonstrating goodness of fit, see Supplemental Table ST2, Shipley 2000). To visualize the effects of interactions, we plotted one predictor against different levels of the interacting predictor, holding all other predictors at their mean (R package ‘*visreg’* version 2.7.0, Patrick Breheny, 2020). This included the following interactions: 1) initial sediment strength and initial crab density to final burrow density; 2) initial *S. alterniflora* live shoot density and final sediment strength to spartina biomass (i.e., *S. alterniflora biomass*); and 3) burrow density to decomposition for each site and depth. Our *‘visreg’* plots draw from the same models constructed for our SEMs for each ecosystem function.

## RESULTS

### Overview of Causal Model Results

Broadly, site, year, and block (i.e., replication) did not affect relationships between predictor and response variables in the linear and generalized linear models we used to build our three Structural Equation Models (SEMs). Although there were differences in magnitude for each site, year, and block, we observed similar trends (i.e., the same slope) in fiddler crab direct and indirect effects of ecosystem functioning (Supplemental Figure SF3A-C, Supplemental Table ST3-ST6). Therefore, the overall effect of fiddler crabs was the same in historical as well as range expanded habitats (with one exception, see *Decomposition* below). For simplicity, we have omitted site, block, and year from path diagrams (except where site is an interacting covariate), but describe site, year, and block effects throughout the results section below.

### Sediment Strength

While more crabs led to more burrows (p=0.009), the number of burrows created per crab increased with increasing sediment hardness (Figures 1, 2, Supplemental Tables ST3, p=0.004, overall model R^2^=0.74). More burrows then led to lower final sediment strength (F_15,80_=4.94, p=0.03, overall model R^2^=0.65), regardless of between site and between year variability. High initial sediment strength increased final burrow density (p=0.03, Supplemental Table ST3), and initial sediment strength significantly influenced final sediment strength (F_15,80_=14.62, p=0.0002, Supplemental Table ST4). The sign of the relationship between initial and final sediment strength differed by site, however. This is demonstrated by the significant interaction term between initial sediment strength and site on final sediment strength (F_15,80_=9.38, p=0.003, Supplemental Table ST4). We ran a linear model on just the relationship between initial and final sediment strength in PIE and NAN separately. In PIE, there was a significant positive relationship between initial and final sediment strength; plots with high initial sediment strength had significantly higher final sediment strength versus plots with low initial sediment strength (F_7,40_=20.44, p<0.0001, Supplemental Table ST5, Supplemental Figure SF4). Alternatively, in NAN the sign flips, and final sediment strength is significantly lower in plots with high initial sediment strength versus plots with low initial sediment strength (F_7,40_=4.06, p=0.05, Supplemental Table ST5, Supplemental Figure SF4). Nevertheless, this site and initial sediment strength interaction is independent of any crab effect, signaling similarity in how fiddler crabs affect sediment strength in both experimental locations, despite inter-site variability.

**Figure 1:**
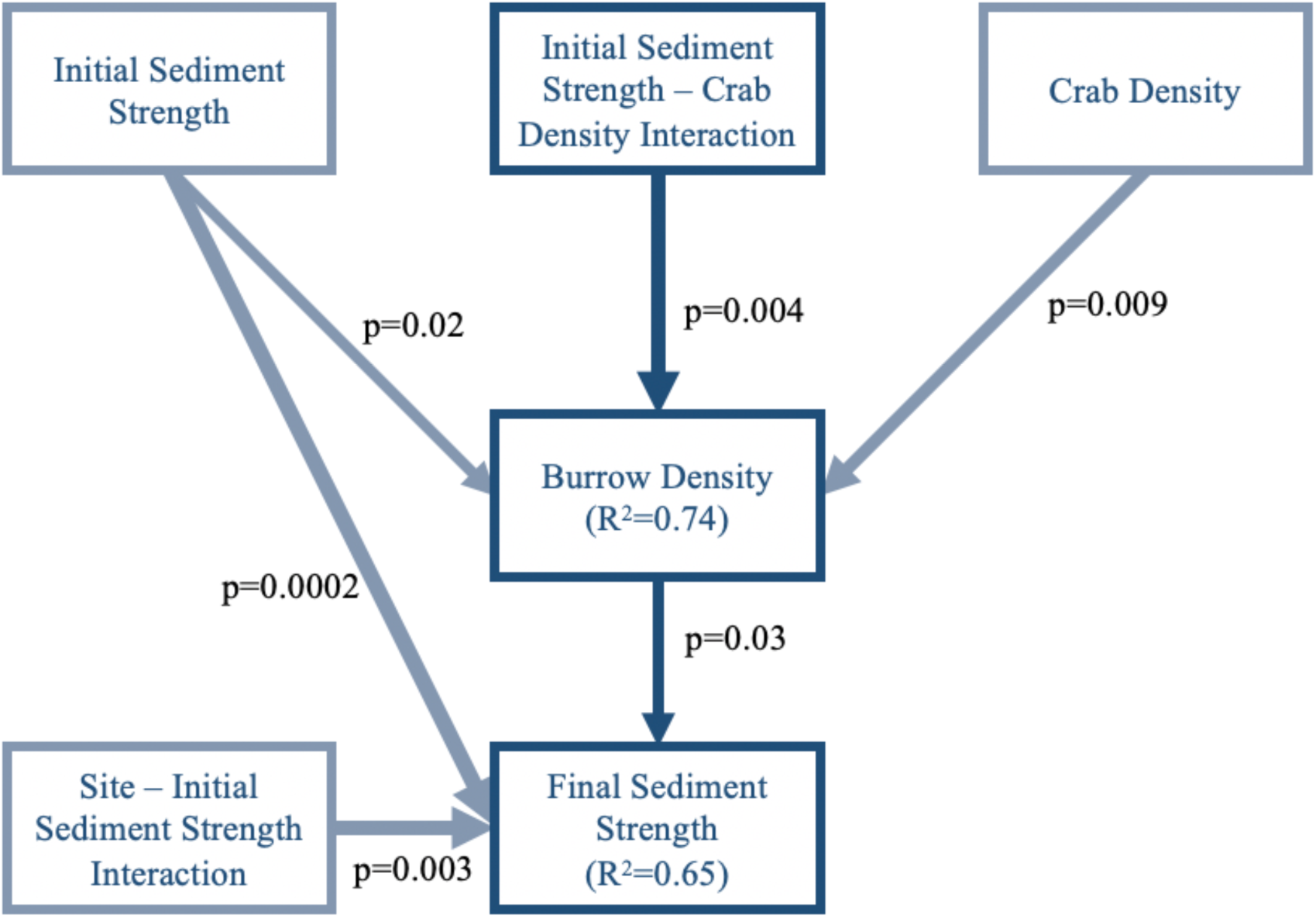
Structural Equation Model (SEM) demonstrating causal connections among: 1) crab density (i.e., fiddler crab density at the beginning of the experiment, crabs 0.25m^−2^); 2) burrow density (i.e., fiddler crab burrow density at the end of the experiment, burrows 0.25m^−2^); 3) initial sediment strength (kPa); 4) final sediment strength (kPa); 5) a site and initial sediment strength interaction; and 6) the crab density-initial sediment strength interaction. Line thickness denotes the degree of significance; thick lines indicate a significant pathway, and thin lines indicate a non-significant pathway (all pathways in this model are significant). Dark blue lines highlight the direct pathway between the crab density initial-sediment strength interaction term and final sediment strength. Thick gray lines indicate significant pathways among the other variables.

**Figure 2:**
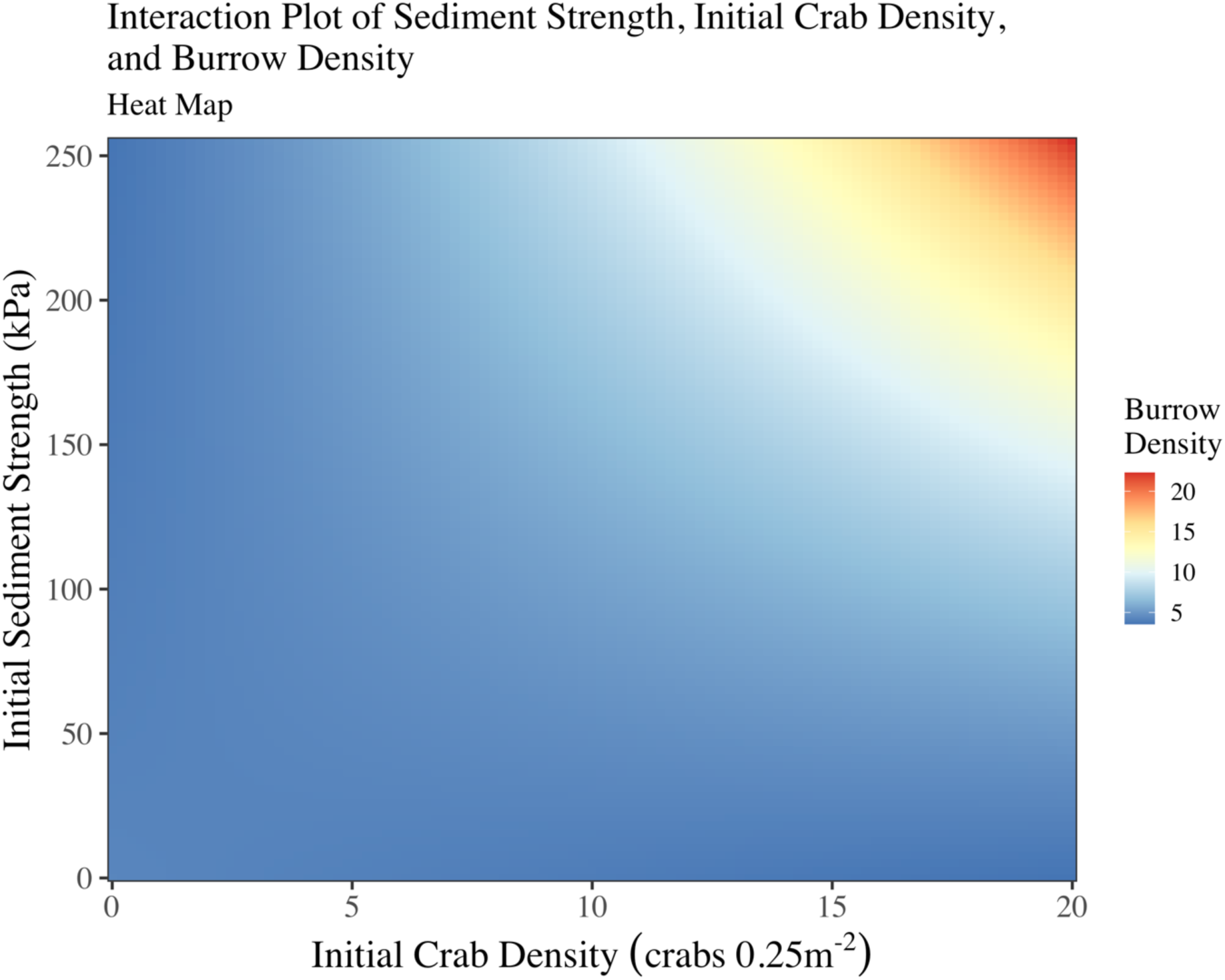
A model generated *visreg* heatmap demonstrating the relationship between initial crab density (crabs 0.25m^−2^, x-axis) and initial sediment strength (kPa, y-axis) with burrow density (burrows 0.25m^−2^, right key). Warmer colors indicate more burrows per plot, cooler colors indicate fewer burrows (i.e., burrow density is highest when both crab density and initial sediment strengths are highest).

Broadly, our SEM also demonstrates significant differences between sites, between years, and among blocks for burrow density, initial sediment strength, and final sediment strength, with a non-significant effect of site on final sediment strength by itself (p=0.07). We found higher burrow densities in NAN (15.0±1.3 burrows plot^−1^) versus PIE (2.5±0.4 burrows plot^−1^, p<0.00001), higher initial sediment strengths in PIE (145.2±8.4kPa) versus NAN (57.9±6.6kPa, F_15,80_ = 81.28, p<0.00001), and blocks differing for each site across all models. We found higher burrow densities in 2018 (12.1±1.6 crabs plot^−1^) versus 2017 (5.4±0.7 crabs plot^−1^, p<0.0001), higher initial sediment strengths in 2018 (119.9±10.2kPa) versus 2017 (83.2±8.8kPa, p=0.007), and block differences across all models.

### Primary Production

Our SEM analysis of fit models for primary production show the effect of fiddler crabs on *Spartina alterniflora* aboveground biomass is indirect. Plots with high *S. alterniflora* shoot density and low sediment strength had the highest *S. alterniflora* aboveground biomass (Figures 3, 4, Supplemental Table ST6, p=0.009, overall model R^2^=0.31) regardless of between site and between year variability. As we showed in our sediment strength SEM, more burrows led to weaker sediment. Additionally, plots with low sediment strength and high *S. alterniflora* shoot density had the highest aboveground *S. alterniflora* biomass. High initial sediment strength led to lower *S. alterniflora* biomass (p=0.0001), and plots with low initial live shoot density and high final live shoot density had high *S. alterniflora* aboveground biomass (i.e., the interaction between initial and final live shoot density, p<0.0001) irrespective of variability between sites and years (Supplemental Figure SF5). High initial live shoot density predicted higher densities of *S. alterniflora* live shoots at the end of the experiment (Figure 3, p=0.0002, overall model R^2^=0.69) for both sites and years. Neither *Minuca pugnax* density (p=0.62), nor *M. pugnax* burrow density (p=0.95) affected final live shoot density of *S. alterniflora*. No interactive effects among predictor variables in our final live shoot density model caused a change in final live shoot densities per plot.

**Figure 3:**
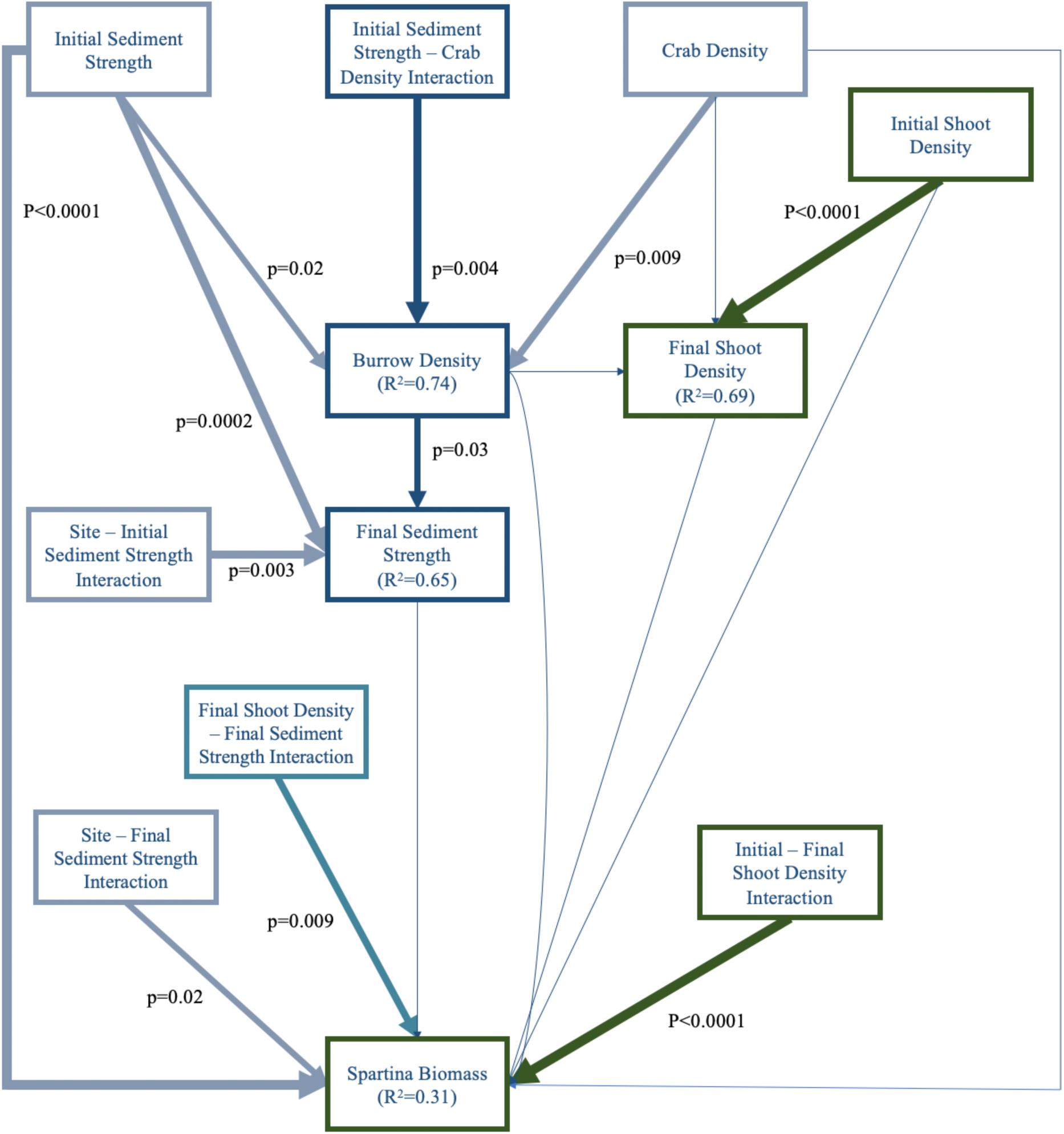
Structural Equation Model (SEM) that incorporates the sediment strength SEM from Figure 1 with indicators of primary production. Line thickness denotes the degree of significance; thick lines indicate a significant pathway, and thin lines indicate a non-significant pathway. Blue lines highlight the direct pathway between the fiddler crab density (crabs 0.25m^−2^) and initial sediment strength (kPa) interaction term and final sediment strength (kPa). Dark green lines indicate pathways related to primary production (i.e., initial and final *Spartina alterniflora* live shoot density, shoots 0.125 m^−2^, and *S. alterniflora* aboveground biomass, g). Teal represents the interaction between final sediment strength and final *S. alterniflora* live shoot density. Thick gray lines indicate significant pathways among all other variables.

**Figure 4:**
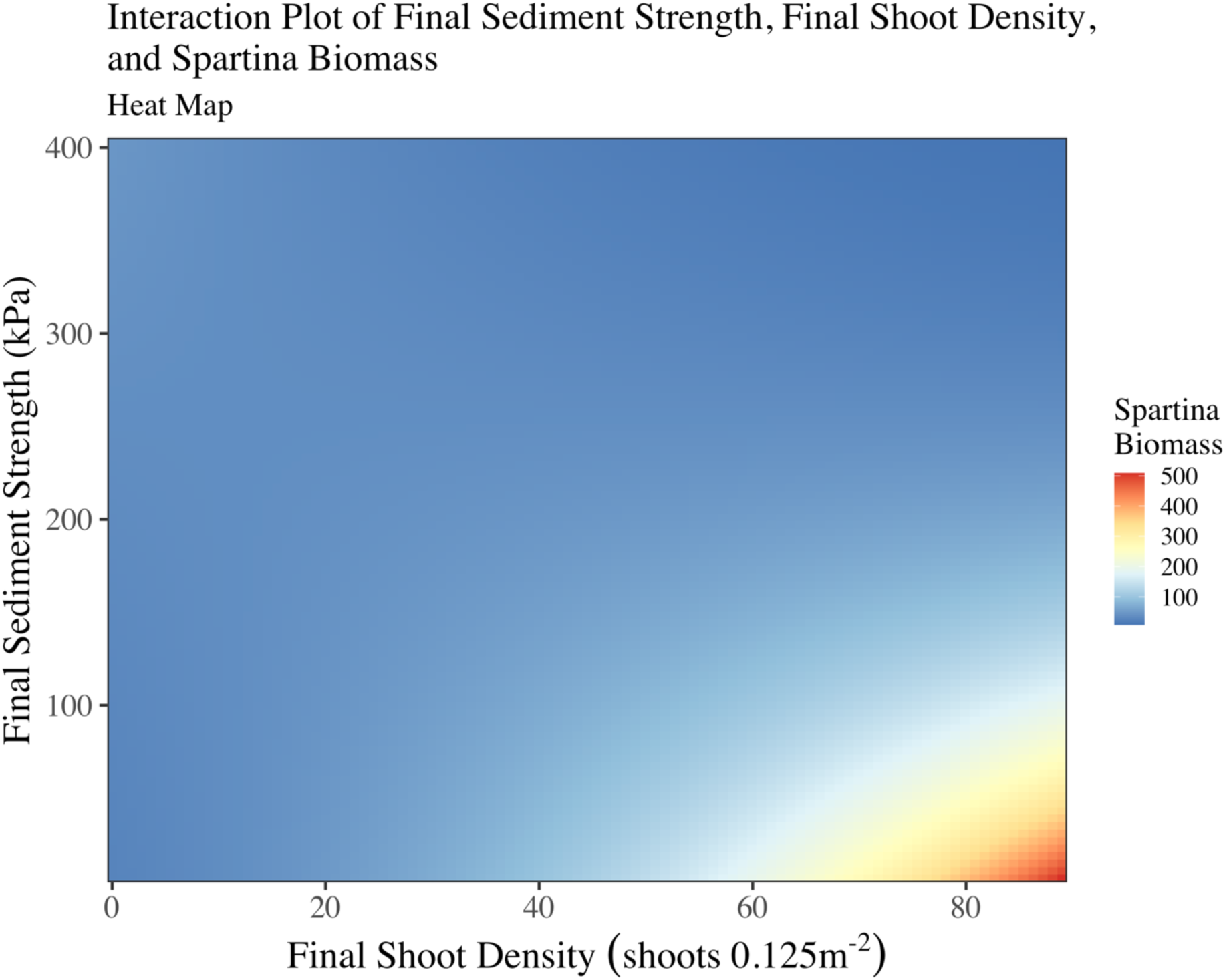
A model generated *visreg* heatmap demonstrating the relationship between final *Spartina alterniflora* live shoot density (shoots 0.125m^−2^, x-axis) and final sediment strength (kPa, y-axis) with *Spartina alterniflora* aboveground biomass (g 0.125m^−2^, right key). Warmer colors indicate more *S. alterniflora* aboveground biomass, cooler colors indicate less biomass (i.e., *S. alterniflora* biomass is highest at high final shoot densities and low final sediment strengths).

Initial and final *Spartina alterniflora* live shoot densities were significantly higher in 2018 (initial: 45.6±2.3 shoots 0.125m^−2^; final: 41.8±2.2 shoots 0.125m^−2^) than 2017 (initial: 19.9±1.8 shoots 0.125m^−2^; final: 12.9±1.0 shoots 0.125m^−2^, both p<0.0001). Initial shoot density was higher in NAN (38.6±2.8 shoots 0.125m^−2^) versus PIE (26.9±2.5 shoots 0.125m^−2^, p=0.004), but final shoot density was not (NAN - 28.4±2.8 shoots 0.125m^−2^; PIE - 26.2±2.6 shoots 0.125m^−2^, p =0.58), and did not differ among blocks (p=0.99). *S. alterniflora* aboveground biomass was higher in PIE (67.8±6.4 g 0.125m^−2^) versus NAN (34.8±2.9 g 0.125m^−2^, p<0.0001), and differed among blocks (p=0.02). *S. alterniflora* aboveground biomass also was higher in 2018 (65.1±6.2 g 0.125m^−2^) versus 2017 (37.5±3.8 g 0.125m^−2^; p<0.0001). In addition, the effect of final sediment strength on *S. alterniflora* aboveground biomass differed by site (p=0.02). In PIE, stronger sediment was correlated with high *S. alterniflora* aboveground biomass; however, in NAN, there is no direct relationship between final sediment strength and aboveground grass biomass (independent of a crab effect). We nevertheless observed the interactive effects of final sediment strength and final *S. alterniflora* live shoot density with *S. alterniflora* biomass described above even after including these site level differences in our generalized linear model for *S. alterniflora* aboveground biomass. The relationship between sediment strength and grass biomass therefore appears to be indirectly driven by fiddler crab burrow density.

### Decomposition

Plots with the highest burrow densities at the shallowest depths (i.e., 5cm versus 15cm below the sediment surface) significantly predicted changes in decomposition broadly (Figure 5, 6, F_15,80_ = 8.45, p=0.005, overall model R^2^=0.46). We found differences in how fiddler crabs manipulate decomposition between sites, however (F_7,40_ = 7.10, p=0.01). In NAN, more burrows at 5 cm yielded the highest decomposition; at PIE however, we saw no crab effect on decomposition at either depth (see Figure 6). In other words, we observed 2 two-way interactions: a depth-burrow interaction on decomposition (p=0.005), and a burrow-site interaction on decomposition (p=0.01). Interestingly, this did not translate to a site-depth-burrow three-way interaction (p=0.60). Initial sediment strength also predicted differences in decomposition, such that plots with higher initial sediment strengths had significantly higher final litterbag decomposition (i.e., lower decomposition) relative to plots with lower initial sediment strength (F_7,40_ = 5.42, p=0.02).

**Figure 5:**
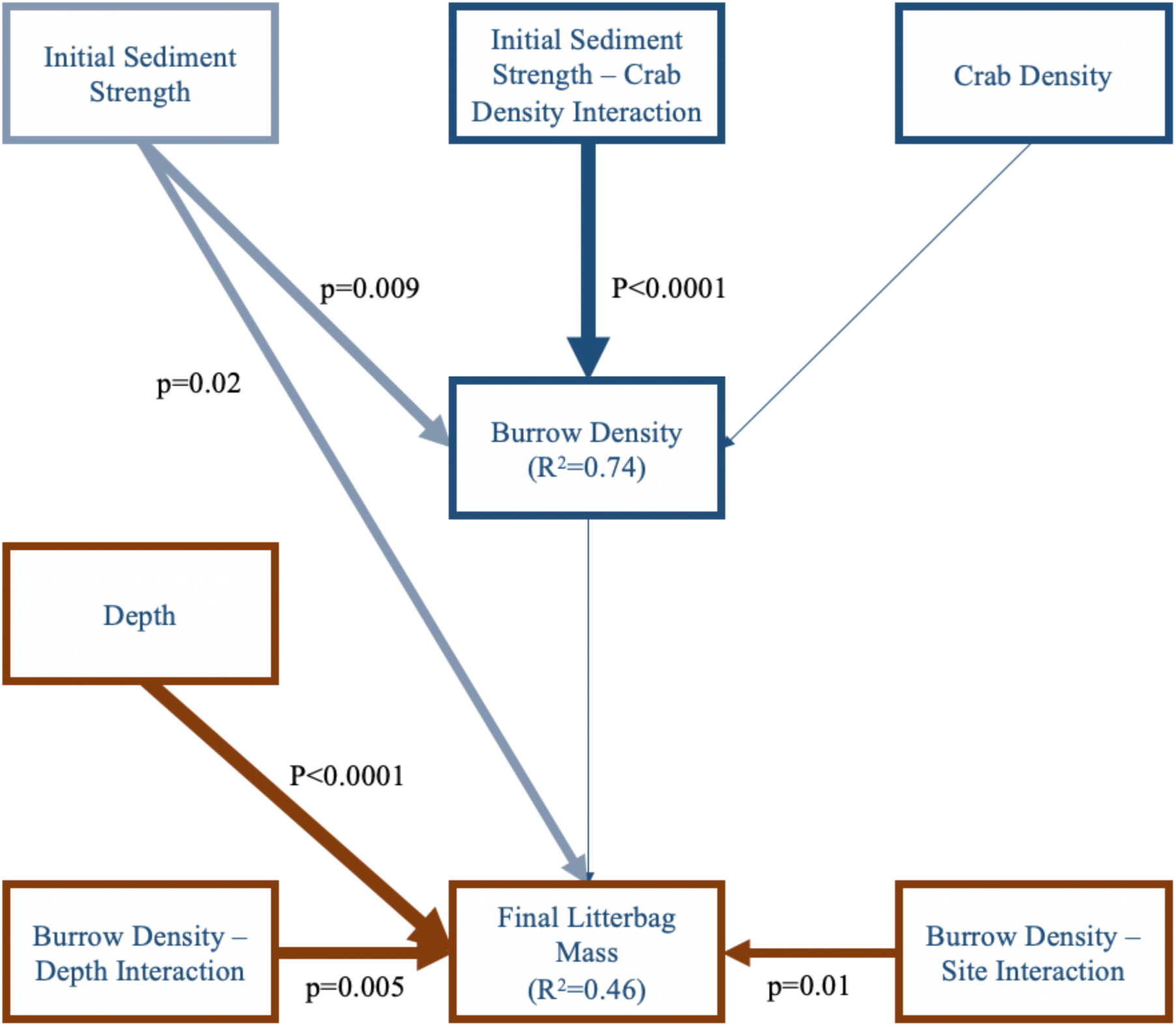
Structural Equation Model (SEM) demonstrating causal connections among 1) crab density (i.e., fiddler crab density at the beginning of the experiment, crabs 0.25m^−2^); 2) burrow density i.e., fiddler crab burrow density at the end of the experiment, burrows 0.25m^−2^); 3) initial sediment strength (kPa); 4) a crab density and initial sediment strength interaction; 5) depth (i.e., litterbag decomposition assay burial depth, cm); 6) a burrow density and depth interaction; 7) a burrow density and site interaction; and 8) final litterbag mass (i.e., decomposition, g). Line thickness describes the degree of significance; thick lines indicate a significant pathway, and thin lines indicate a non-significant pathway. The blue line highlights the direct pathway between the crab density and initial sediment strength interaction term with final sediment strength. The thick gray line indicates a significant pathway between initial sediment strength and final litterbag mass. Brown lines and boxes specify linkages for the direct decomposition components to the SEM. Note, two non-significant site interactions were omitted from this diagram for clarity: a site and depth two-way interaction (p=0.08) and a depth, burrow density, and site three-way interaction (p=0.60).

**Figure 6:**
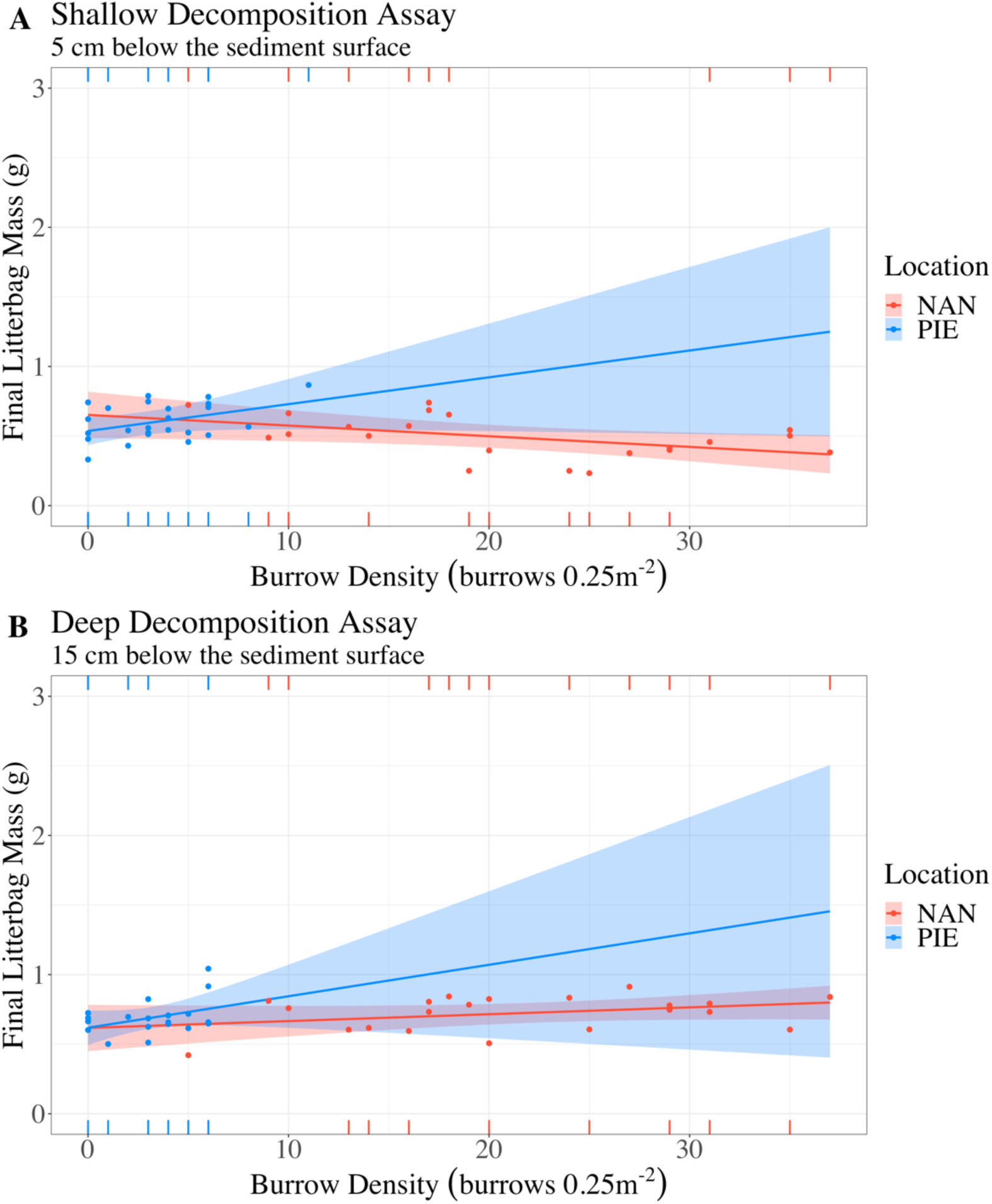
**A** (top panel) the relationship between crab burrow density (burrows plot^−1^, x-axis) and decomposition (i.e., final litterbag mass, g) for each site NAN (i.e., Nantucket) and PIE (i.e., Plum Island Estuary) for litterbags buried at 5cm below the sediment surface (i.e., shallow decomposition assay). **B** (bottom panel) the same relationships but for litterbags buried at 15cm below the sediment surface (i.e., deep decomposition assay). Plots are model generated *visreg* linear regressions. Note the wide confidence intervals for PIE indicating unknown effects of burrows on decomposition at high fiddler crab burrow densities.

There was no significant difference in mean decomposition between sites (higher final mass = lower decomposition – NAN: 0.66±0.03g litterbag^−1^; PIE: 0.60±0.02g litterbag^−1^, p=0.1), but there was a significant site-block interactive effect on decomposition (F_7,40_ = 4.06, p=0.05). Finally, decomposition decreased with increasing depth for both NAN (5cm below sediment surface: 0.57±0.04g litterbag^−1^; 15cm below sediment surface: 0.75±0.03g litterbag^−1^) and PIE (5cm below sediment surface: 0.55±0.03g litterbag^−1^; 15cm below sediment surface: 0.66±0.03g litterbag^−1^, F_7,40_ = 23.77, p<0.0001). As noted, we only used data for 2018 in this model, which qualitatively did not alter the significant relationships observed for the sediment strength and primary production SEMs, with one exception. Fiddler crab density by itself had no effect on the number of burrows at the end of the experiment (i.e., burrow density, p=0.10). As with the sediment strength and primary production SEMs, the interaction between initial sediment strength and fiddler crab density caused higher burrow densities, however (p=0.00007). Finally, we lost 13 out of 112 litterbags (~12%) below the sediment surface (11 from caged plots, 2 from uncaged control plots), unbalancing replications and potentially biasing our decomposition results. Nevertheless, our experimental design is relatively robust to missing data (i.e., establishing 6 density treatments replicated 4 times per site per year, Cottingham et al. 2005), and our tests of model fit for decomposition resulted in no violations of assumptions (see Supplemental Methods 2). Therefore, we feel confident in the results we present here for the effect of fiddler crabs on decomposition.

## DISCUSSION

Our results demonstrate that local environment can modify the impact of a range-expanding ecosystem engineer in both historical and expanded habitats. In this case, substrate hardness increased the ability of *Minuca pugnax* (=*Uca pugnax*) to form burrows, which in turn amplified its impact on multiple ecosystem functions. More burrows meant a greater decline in sediment strength. Burrowing activity that weakened sediment was also associated with increased aboveground *Spartina alterniflora* biomass. Fiddler crabs increased decomposition at shallow (i.e., 5cm) but not deep (i.e., 15cm) depths below the sediment surface in Nantucket, regardless of burrow density. We did not observe the same effect on decomposition in Plum Island, potentially due to the lower relative density of burrows in PIE versus NAN by the end of the experiment (up to 5 times more burrows in NAN than in PIE). Unlike sediment strength and primary production, a critical threshold of burrows may be required to influence decomposition (i.e., increased soil oxygenation and soil microbe activation by higher burrow densities, Gribsholt et al. 2003). In each case, however, fiddler crabs affected historical and expanded habitats through a series of direct and indirect interactions. Taken together, our results demonstrate that a range expanding ecosystem engineer is impacted by and impacts its expanded and historical ranges via similar mechanisms, in this case related to sediment stability and primary production.

We make these inferences based on our causal pathways of influence from fiddler crabs to ecosystem properties. The precise mechanisms behind these pathways remain unclear in some cases, although the natural history of the system provides some clues. The observed decline in sediment strength due to mud fiddler crab burrowing activity could have been the result of increased saturation of marsh sediment or decreased belowground biomass of *Spartina alterniflora*, or their combination. For salt marshes along the east coast of the United States, *M. pugnax* both decreases root biomass and increases *S. alterniflora* aboveground biomass (Bertness 1985, Gittman and Keller 2013). The mud fiddler crab also increases redox potential via their burrows, which increases available nutrients for plant root uptake (Thomas and Blum 2010, Gittman and Keller 2013). Several studies demonstrate that increased nutrient availability decreases *S. alterniflora* belowground biomass, thereby decreasing both root mat density and overall sediment strength (Deegan et al. 2012, Jafari et al. 2019, Crosby et al. 2021). Our study supports these results; plots with the highest aboveground biomass had the lowest sediment strength and the highest *S. alterniflora* live shoot density. Altogether, there is support for fiddler crabs increasing available nutrients in our plots, facilitating both increased aboveground biomass and reduced belowground biomass in both study sites. Future research should mechanistically link burrows with sediment strength, dissolved nutrients, and above and belowground *S. alterniflora* biomass in both historical and expanded New England salt marsh habitats.

Additionally, we observed the same effects of fiddler crab burrows on functioning in both locations despite higher burrow densities in NAN by the end of the experiment. Therefore, the *per capita* effect of fiddler crabs and their burrows in PIE was greater than NAN. Martínez-Soto and Johnson (2020) also showed significantly lower fiddler crab densities in PIE relative to their historical range, which is common for range-expanding species. Species at the beginning of their expansion (like *M. pugnax*) can experience barriers to establishing dense populations such as settlement dynamics and allee effects (Keith et al. 2011, Walter et al. 2017). However, we observed significantly lower burrow densities in PIE relative to Nantucket despite keeping initial fiddler crab density the same between locations for both years. There is likely some other barrier to maintain densities comparable to southern, historical habitats; however, PIE crabs could be compensating for this via selection over time. Wong et al. (2021) demonstrated that, even controlling for size differences, PIE crabs are better at burrowing in high sediment strengths than Nantucket crabs. If fiddler crab burrow densities in PIE reach levels as high as their historical ranges over time, they could have stronger population level effects in expanded salt marsh habitats than historical ones for some functions (e.g., sediment stability and primary production), but not others (e.g., decomposition).

Sea level rise and marsh edge erosion in PIE and elsewhere could be impacted by the presence of a recently established range-expanding ecosystem engineer (Gedan et al. 2011, Wigand et al. 2017). Our research demonstrates that crabs weaken sediment strength, which could exacerbate climate induced marsh instability. We also found that fiddler crabs need both strong sediment and high crab densities to facilitate large burrow densities, however. As fiddler crabs increase in density in expanded ranges, their capacity to burrow reduces with increased crab densities, potentially facilitating a balancing feedback loop controlling their population. Fiddler crabs, in both historical and expanded ranges, can also influence *S. alterniflora* seed dispersal, control algal densities, influence bioturbation, and increase sequestered carbon, which all are involved in marsh stabilization (Gutiérrez et al. 2006, Thomas and Blum 2010, Smith and Tyrrell 2012, Smith and Green 2015, Johnson et al. 2020). Additionally, fiddler crabs could be indirectly facilitating increased capture of suspended sediments in the incoming tides by enhancing aboveground plant biomass observed in our study (Kirwan et al. 2009, Morris et al. 2013). Roy et al. (2023) showed that simulated marsh growth enhancement by fiddler crabs effectively mitigated losses in marsh area due to high sea level rise over time (i.e., greater than 18mm year^−1^ by the year 2100). Crabs may facilitate relative marsh stabilization over long time horizons, especially if their densities reach levels comparable to historical habitats.

Our study provides critical context for researchers studying range expansions both as an individual process and as part of a suite of processes related to environmental change. Range expansions ecology should therefore focus on a more thorough understanding of the mechanistic links between range-expanding species and their surrounding historical ecosystems, particularly for ecosystem engineers such as fiddler crabs. Ecosystem engineers globally are shifting their distributions, and could be influencing sediment strength, nutrient availability, and primary production in their new residences similar to *M. pugnax* (Edgar et al. 2005, Sorte et al. 2010, Balvanera et al. 2014, Hu and Juan 2014). Yet, we do not fully understand how species in expanded habitats affect ecosystem functioning *relative to* historical habitats. Invasive species research does not provide sufficient insight into native range expansion dynamics due to differences inherent between bioinvasions and range expansions (Sorte et al. 2010, Wallingford et al. 2020). Our study provides evidence that untangles such dynamics in range expansions ecology. We found similarity in how *M. pugnax* influences ecosystem functioning in its natal versus expanded range. The impact of the mud fiddler crab in its expanded range is predictable from their historical habitats, contrary to our expectations based on differences in site characteristics. Importantly, tests of conditional independence or explicit tests of direct effects do not indicate that we have any missing paths from fiddler crab population and burrow density to these functions. Their impact was entirely affected by burrow density. We therefore emphasize the need for researchers to study relative impacts of range expanders between expanded and historical habitats globally, and utilize causal mechanisms of statistical inference (i.e., SEM) as demonstrated here.

Species range expansions due to climate change are becoming more frequent. Our study demonstrates consistency between historical and expanded regions and represents one of the first key insights into how an ecosystem engineering range-expander will interact with a habitat in its new range *relative to* its historical habitat. This research also shows that physical properties directly influence crab burrowing behavior in the same ways in both historical (i.e., Nantucket) and expanded (i.e., PIE) habitats. *M. pugnax* in turn indirectly affects sediment strength of both ranges via burrowing, causing cascading effects that influence primary production (in both habitats) and decomposition (at shallow depths in its historical habitat). This system provides researchers with the ability to better understand how range-expanders influence current and future ecosystem functioning in expanded habitats globally based on ecological knowledge in historical ranges. Representing novel understanding for this species and the field of range expansions overall, this study provides critical insight into how range-expanders may be influenced by, while simultaneously influencing, ecosystem structure and function in historical versus expanded ranges.

## Supporting information

2024_Roy_Johnson_Byrnes_Supplement

## ACKNOWLEDGEMENTS

This work was supported by Anne Giblin and the National Science Foundation as part of the Plum Island Estuary Long Term Ecological Research (PIE-LTER) program (no. 1637630), Yvonne Vaillancourt and the Battle Fund at the University of Massachusetts Boston Nantucket Field Station, the Sanofi-Genzyme Dean’s Fellowship and Beacon Student Success Fellowship at the University of Massachusetts Boston, and the National Science Foundation Coasts and Communities Integrated Graduate Education and Research Traineeship (NSF IGERT). We thank Barbara Araya-Naranjo, Alex Walus, Mark Hensel, Maeve Forbes, Charlotte Whittier, Linnea Sturdy, and Richard Wong for their invaluable assistance in this project. This is contribution XXXX from the Virginia Institute of Marine Science, William & Mary.

There are no conflicts of interest to disclose.

## AUTHOR CONTRIBUTIONS

Michael S. Roy, David Samuel Johnson, and Jarrett E. K. Byrnes conceived the overall research ideas; Michael S. Roy and Jarrett E. K. Byrnes analyzed the data; Michael S. Roy led the writing of the manuscript and collected the data; Michael S. Roy, David Samuel Johnson, and Jarrett E. K. Byrnes contributed to critical review, commentary, and revision of the manuscript; Michael S. Roy, David Samuel Johnson, and Jarrett E. K. Byrnes developed and designed the methodology; Michael S. Roy developed and created the visualizations. All Authors contributed critically to the drafts and gave final approval for the publication.

