## Supplementary material for "A range expanding ecosystem engineer influences historical and expanded habitats via the same causal pathways": 2024_Roy_Johnson_Byrnes_Supplement

#### Supplemental Material

#### SUPPLEMENTAL METHODS 1

The UMass Boston Nantucket Field Station is located on the north side of the island of Nantucket, Massachusetts (i.e., NAN). Encompassing 103 acres, the field station contains a diversity of habitats in addition to salt marsh that include sand dunes, eelgrass beds, and terrestrial scrub brush. Being south of Cape Cod, MA, air and water temperatures of Nantucket are strongly influenced by the gulf stream (Gawarkiewicz, Todd, Plueddemann, Andres, & Manning, 2012). As a result, Nantucket is species rich relative to habitats north of the Cape, supporting myriad species including the purple marsh crab *Sesarma reticulatum,* the blue crab *Callinectes sapidus*, and the mud fiddler crab *Minuca pugnax* (the focus of this study, Pennings and Bertness 2001). *M. pugnax* burrows along NAN creek banks in the lower elevation marsh zone dominated by tall *Spartina alterniflora*, encountering a tidal range of ~1m (Luk & Zajac, 2013). As a burrowing species of crab, sediment hardness could play a significant role in the survival of this species, which is on average ~10psi (~69 kPa) in NAN due primarily to silty oceanic inputs of sediment (*pers. obs*, Langlois 1979).

At our study site north of the Cape, the Plum Island Estuary (PIE) is the largest continuous salt marsh in the northeastern United States (Langston, Durán Vinent, Herbert, & Kirwan, 2020). It ranges from Gloucester, MA in the south to Seabrook, NH in the north, approximately 60km$^2$ in total. Plum Island is a barrier island that faces the open north Atlantic to the east, with 40km$^2$ of salt marsh to the west. PIE salt marshes of the GoM are on average colder than marshes south of Cape Cod MA due to the absence of the influence of the Gulf Stream and the presence of the cold Labrador Current; however, the GoM has experienced relatively rapid warming over the least 40 years (Gonçalves Neto, Langan, & Palter, 2021). PIE's tidal range is ~3m between high and low tides with a mean sediment strength at ~25-50psi (~172-345 kPa) due to primarily peaty organic rich sediment from low turnover of decomposing material and riverine inputs of sediment (Deegan et al., 2012; Johnson, Warren, Deegan, & Mozdzer, 2016; Vincent, Burdick, & Dionne, 2013). As with NAN, fiddler crabs burrow in the *Spartina alterniflora* dominated lower elevation zone of the salt marsh in PIE (Martínez-Soto & Johnson, 2020).

**Supplemental Table ST1: Control Plot Comparisons**

| Predictor | Response | Estimate | Std. Error | t-Value | p-Value | Site |
|---|---|---|---|---|---|---|
| Crab Density | Initial Shoot Density | 2.375 | 10.87 | 0.22 | 0.83 | NAN |
| Crab Density | Initial Sediment Strength | 1.208 | 3.16 | 0.38 | 0.708 | NAN |
| Crab Density | Final Shoot Density | 1.75 | 10.71 | 0.16 | 0.873 | NAN |
| Crab Density | Final Sediment Strength | -0.773 | 3.69 | -0.21 | 0.837 | NAN |
| Crab Density | S. alterniflora Aboveground Biomass | 12.963 | 11.1 | 1.17 | 0.262 | NAN |
| Crab Density | Initial Shoot Density | -0.375 | 7.1 | -0.05 | 0.959 | PIE |
| Crab Density | Initial Sediment Strength | 0.065 | 3.83 | 0.02 | 0.987 | PIE |
| Crab Density | Final Shoot Density | 8.25 | 7.86 | 1.05 | 0.312 | PIE |
| Crab Density | Final Sediment Strength | 0.73 | 4.65 | 0.16 | 0.878 | PIE |
| Crab Density | S. alterniflora Aboveground Biomass | 7.234 | 26.47 | 0.27 | 0.789 | PIE |

**Table ST1:** For NAN – Nantucket: Test statistics for linear model comparisons with Gaussian error and an identity link between control plot (i.e., no cage) and the twenty-crab cage (i.e., cage with ambient crab densities in Nantucket, Aspey 1978); For PIE – Plum Island Estuary: Test statistics for linear model comparisons between control plot (i.e., no cage) and four-crab plot (i.e., cage with ambient densities in Plum Island Estuary, Martinez-Soto and Johnson 2020). Note, decomposition is not included in this analysis because of a lack of statistical power and an inability to generate unbiased p-values: we only have decomposition data for 2018, and there were 13 out of 112 missing litterbags making the comparisons between plot types unbalanced.

**Supplemental Figure SF1:**

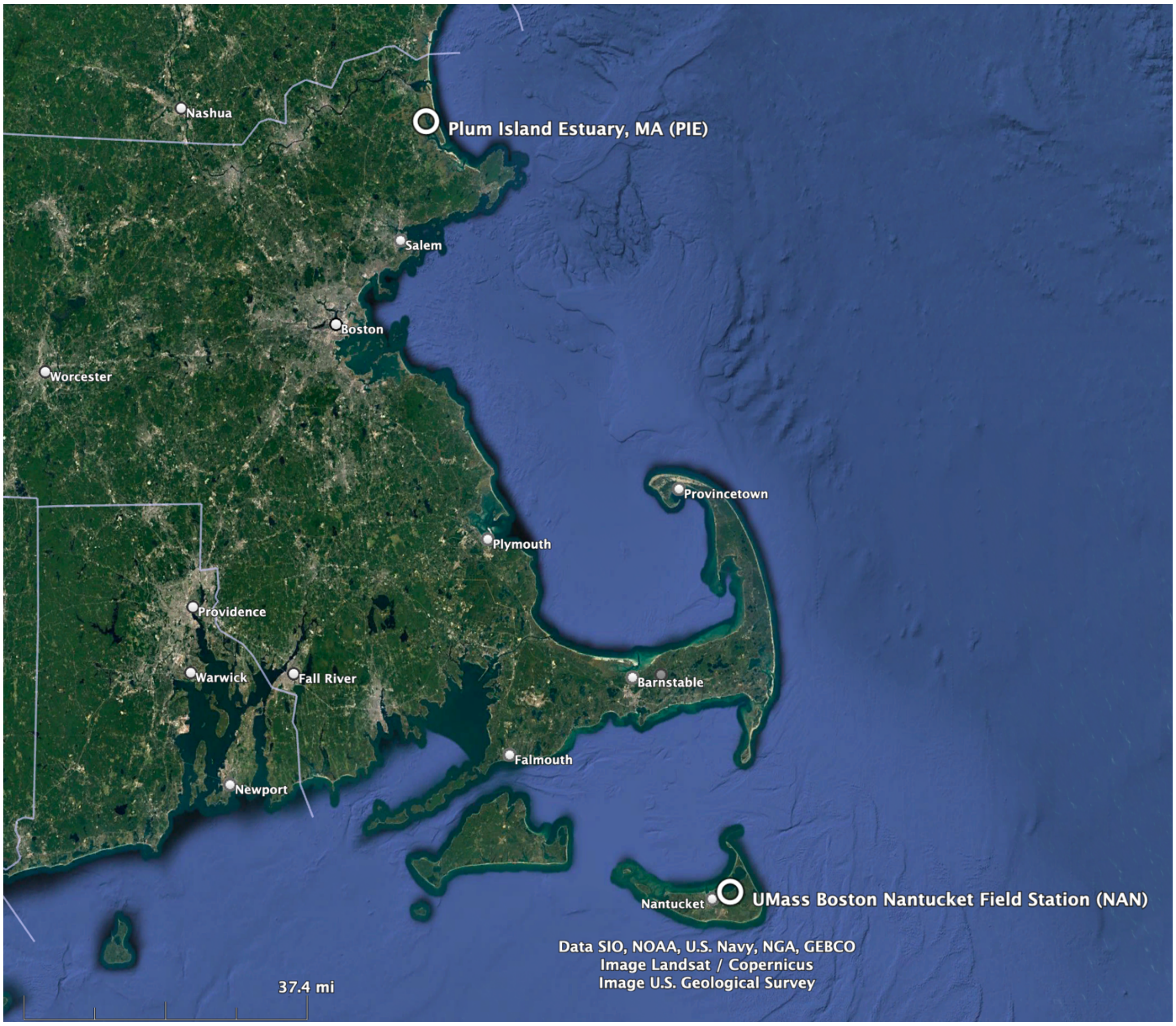


**Figure SF1: Map of Coastal Rhode Island (RI), Massachusetts (MA), and New Hampshire (NH). The Plum Island Estuary (PIE) is in northern MA at the top of the map. The University of Massachusetts (i.e., UMass) Boston Nantucket Field Station (NAN) is on Nantucket Island in southern MA at the bottom of the map, more than 170km from PIE. Map generated from Google Earth.**

**Supplemental Figure SF2: Sediment Strength Measurements Between Years**

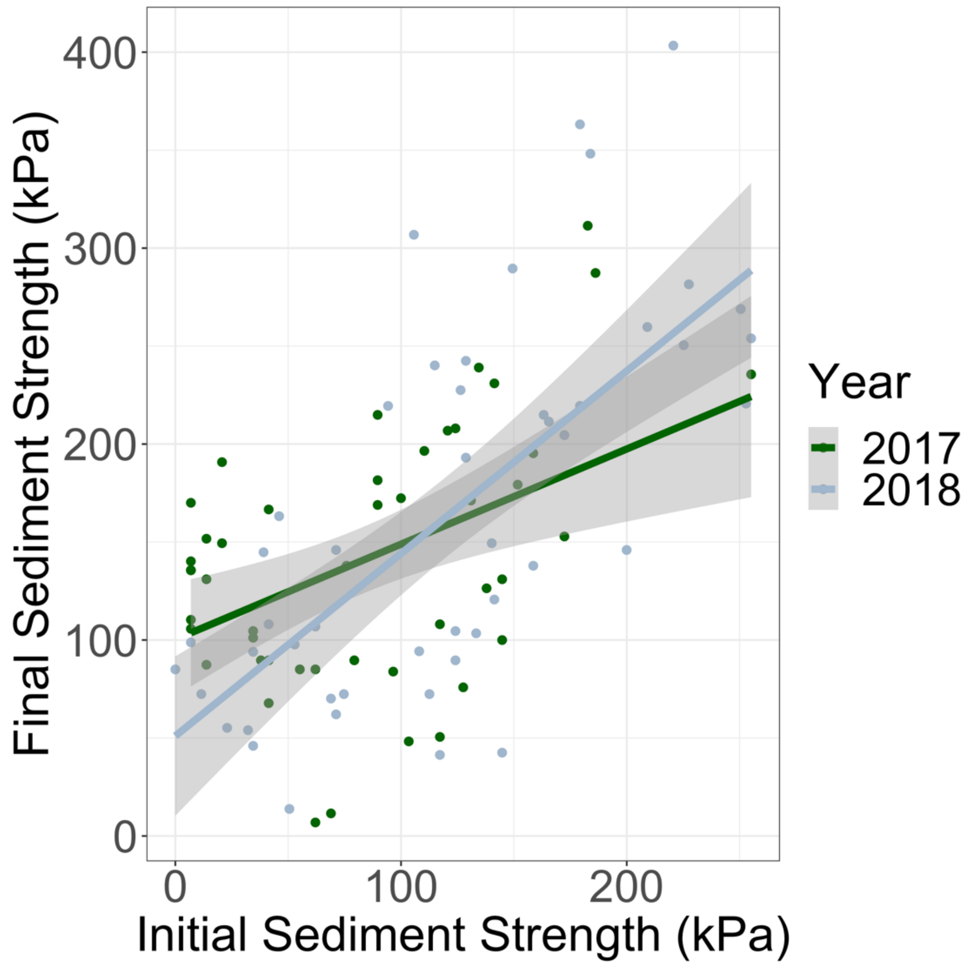


**Figure SF2:** Comparison plot (*ggplot2*) showing no significant difference in slope (i.e., overlapping confidence intervals) between 2017 (green line) and 2018 (grey line) for the relationship between initial (x-axis) and final (y-axis) sediment strength (measured in kilopascals, kPa).

### SUPPLEMENTAL METHODS 2

We tested assumptions for each model using '*fitdistrplus*' (Wotherspoon, Melbourne-Thomas, & Raymond, 2022), and *'car'* (Fox et al., 2016), and '*DHARMa*' (Hartig, 2019) packages in R. No model violated assumptions for each distribution with one exception: our *Spartina alterniflora* aboveground biomass model. First, this model demonstrated a high Variance Inflation Factor (VIF) using the *'vif'* function in the *'car'* package. To correct for high VIFs, we centered our data by taking each data point for each predictor we measured (which did not include initial crab density as that was the parameter we manipulated) and subtracting each point from the mean value of that parameter (Robinson & Schumacker, 2009). After centering our data, VIFs for all parameters shrank below 10 with no change in relationships between predictor and response variables. Therefore, we present uncentered data in our results.

Additionally, in our *Spartina alterniflora* biomass gamma glm, there was a significant deviation in the Kolmogorov-Smirnov (KS) test ($p<0.05$) using the *'simulateResiduals'* function in the '*DHARMa*' package, indicating poor model fit (Lopes, Reid, & Hobson, 2007). Therefore, we re-evaluated our QQ plot of residuals to determine if there were any data points driving this failure in our test of assumptions. Three plots had zero biomass at the end of the experiment in the standardized corner of the plot where we clipped grass. The gamma distribution does not behave well with zeros in the response variable, so we included a correction value of 0.0001g of grass biomass as a proxy for zero (Aksoy, 2000). In fact, for two of the three plots, there were *S. alterniflora* live shoots in the plot, but not in our standardized area where we clipped grass. Therefore, we feel confident that our correction factor accurately depicts *S. alterniflora* biomass within our plots. After adding the correction factor, our model still failed the KS test of assumptions. Therefore, to determine if these three plots were driving the failure in our diagnostic tests of assumptions, we removed them, reran our models, and ran the test again. Removing those values resulted in a non-significant deviation in the KS test ($p>0.05$), indicating a good model fit for the gamma distribution. Therefore, we believe that our statistical model accurately demonstrates the relationships between our predictor and response variables in this model, despite including the three plots with low biomass.

Lastly, we tested whether year interacted with any of our predictors for both our sediment strength and primary production SEM's. We observed one relationship, the interaction between year and initial fiddler crab density significantly affected burrow density ($p=0.03$). Including this interaction did not change the significant relationships we outline in our results section, however. In addition, calculating Akaike Information Criterion (AIC) values for models with and without the year-initial crab density interaction resulted in a $\Delta$AIC (change in AIC) of 2.44, which indicates both models fit similarly to the data (Burnham & Anderson, 2002). Therefore, we omitted this interaction in our burrow density model for clarity.

**Supplemental Table ST2: SEM Goodness of Fit**

| Structural Equation Model | Fisher's C | Degrees of Freedom | p-Value (Goodness of Fit) |
|---|---|---|---|
| Sediment Strength SEM | 2.659 | 4 | 0.62 |
| Primary Production SEM | 12.397 | 14 | 0.57 |
| Decomposition SEM | 6.141 | 8 | 0.63 |

**Table ST2:** Global Goodness of Fit for each ecosystem function structural equation model (SEM). A non-significant p-value indicates there is no significant differences between observed and model generated variance-covariance matrices (i.e., meets the criteria for good model fitting in SEM, Shipley 2000).

#### Supplemental Figure SF3: Comparisons Among Site, Year, and Block

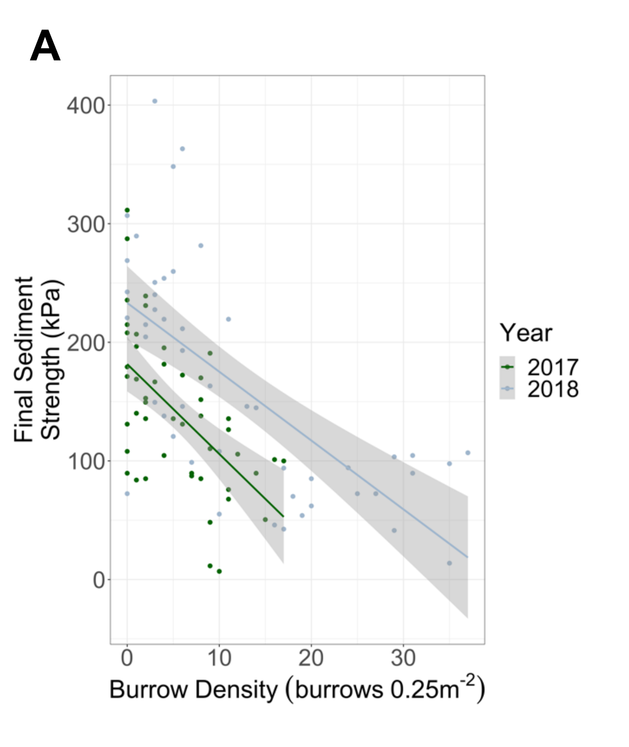


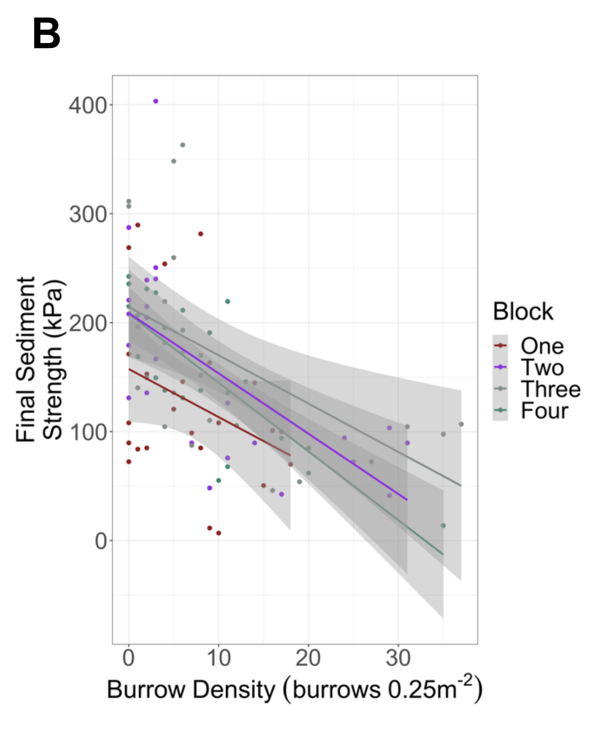


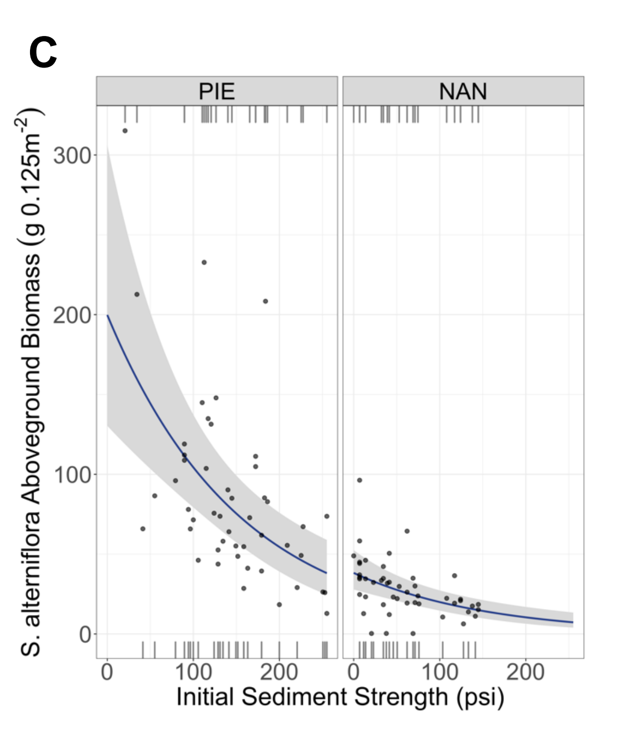


**Figure SF3: A** – Comparison plot (*ggplot2*) between burrow density at the end of the experiment (burrows $0.25m^{-2}$ x-axis) and final sediment strength (kPa, y-axis) between experimental years across sites and blocks (2017 – green, 2018 – gray);
**B** – Comparison plot (*ggplot2*) between burrow density at the end of the experiment (burrows $0.25m^{-2}$ x-axis) and final sediment strength (kPa, y-axis) for each block across sites and years (One – green, Two – Red, Three – Gray, Four - Purple);
**C** – Model generated *visreg* plot between initial sediment strength (kPa, x-axis) and *Spartina alterniflora* aboveground biomass measured at the end of the experiment (g $0.125m^2$, x-axis) between experimental sites PIE (left pane) and NAN (right pane). Demonstrating across metrics differences in scale but not direction (i.e., slope).

**Supplemental Table ST3: Burrow Density Model Test Statistics**

| Predictor | Response | LR Chisq | Df | p-Value |
|---|---|---|---|---|
| Initial Crab Density | Burrow Density | 6.84 | 1 | 0.0089 |
| Initial Sediment Strength | Burrow Density | 5.37 | 1 | 0.0204 |
| Year | Burrow Density | 39.48 | 1 | 0 |
| Site | Burrow Density | 169.92 | 1 | 0 |
| Block | Burrow Density | 5.49 | 1 | 0.0192 |
| Crab Density-Sediment Strength Interaction | Burrow Density | 8.47 | 1 | 0.0036 |

**Table ST3:** Test statistics for a multivariate generalized linear model with a negative binomial error and log link comparing initial crab density (crabs $0.25m^{-1}$), initial sediment strength (kPa), year, site, block, and a 2-way interaction term between initial crab density (crabs $0.25m^{-2}$) and initial sediment strength (kPa) to burrow density (burrows $0.25m^{-2}$). This model was evaluated for significance using Likelihood-Ratio (LR) chi-squared, Type II sums of squares.

**Supplemental Table ST4: Initial and Final Sediment Strength Test Statistics**

| Predictor | Response | Sums Sq | Df | F Value | p-Value |
|---|---|---|---|---|---|
| Year | Initial Sediment Strength | 32454.92 | 1 | 14.34 | 0.0003 |
| Site | Initial Sediment Strength | 183057.8 | 1 | 80.89 | 0 |
| Block | Initial Sediment Strength | 16275.82 | 1 | 7.19 | 0.0087 |
| Burrow Density | Final Sediment Strength | 12490.51 | 1 | 4.94 | 0.0288 |
| Initial Sediment Strength | Final Sediment Strength | 36991.76 | 1 | 14.62 | 0.0002 |
| Year | Final Sediment Strength | 5216.28 | 1 | 2.06 | 0.1546 |
| Block | Final Sediment Strength | 16362.88 | 1 | 6.47 | 0.0127 |
| Site | Final Sediment Strength | 8576.27 | 1 | 3.39 | 0.0689 |
| Initial Sediment Strength-Site Interaction | Final Sediment Strength | 23720.06 | 1 | 9.37 | 0.0029 |

**Table ST4:** Test statistics for a multivariate linear model with Gaussian error and an identity link comparing year, site, and block to initial sediment strength (kPa) and a linear model comparing burrow density (burrows $0.25m^{-2}$), initial sediment strength (kPa), year, block, site, and a 2-way interaction term between initial sediment strength (kPa) and site to final sediment strength (kPa). These two models were evaluated for significance using Type II sums of squares F-test.

**Supplemental Table ST5: Initial to Final Sediment Strength Comparison Between Sites**

| Site | Predictor | Response | Sums Sq | Df | F Value | p-Value |
|---|---|---|---|---|---|---|
| NAN | Initial Sediment Strength | Final Sediment Strength | 6597.39 | 1 | 4.06 | 0.0498 |
| PIE | Initial Sediment Strength | Final Sediment Strength | 77673.36 | 1 | 20.44 | 0 |

**Table ST5:** Test statistics for a univariate linear model with Gaussian error and an identity link comparing initial sediment strength (kPa) to final sediment strength (kPa) for each experimental site (Nantucket – NAN, and Plum Island Estuary – PIE). These two models were evaluated for significance using Type II sums of squares F-test.

**Supplemental Figure SF4: Initial to Final Sediment Strength Comparison Between Sites**

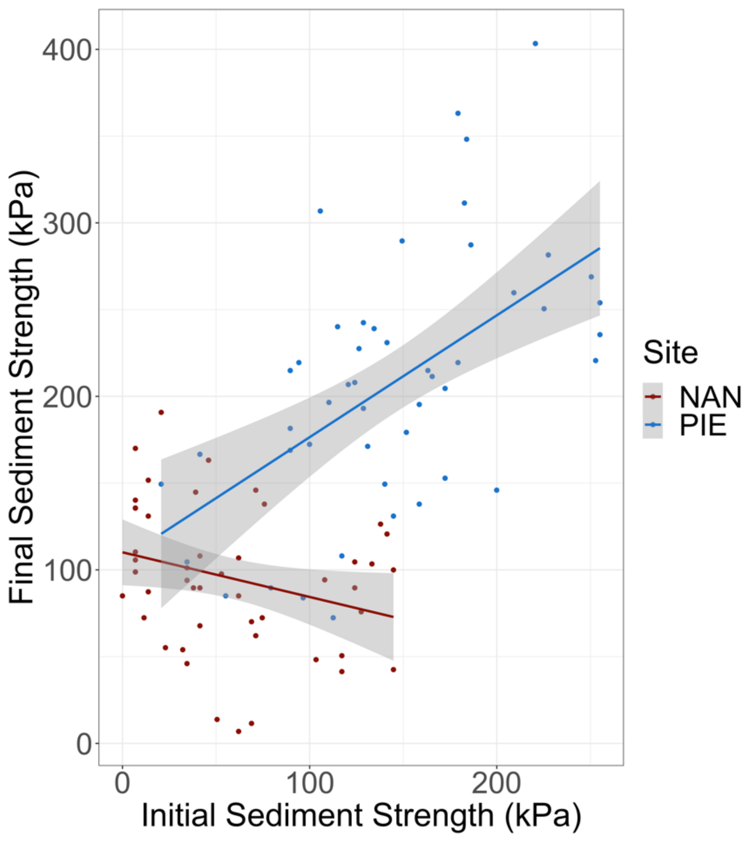


**Figure SF4:** Data generated plot (*ggplot2,* Wickham et al. 2021)*)* showing the relationship between initial sediment strength (kPa) and final sediment strength (kPa) between each site (NAN – Nantucket, red; PIE – Plum Island Estuary, blue).

**Supplemental Table ST6: Initial and Final Spartina alterniflora Shoot Density Test Stats.**

| Predictor | Response | LR Chisq | Df | p-Value |
|---|---|---|---|---|
| Year | Initial S. alterniflora Shoot Density | 98.06 | 1 | 0 |
| Site | Initial S. alterniflora Shoot Density | 26.93 | 1 | 0 |
| Block | Initial S. alterniflora Shoot Density | 0.32 | 1 | 0.5713 |
| Initial Crab Density | Final S. alterniflora Shoot Density | 0.25 | 1 | 0.6177 |
| Initial Sediment Strength | Final S. alterniflora Shoot Density | 2.7 | 1 | 0.1004 |
| Initial Shoot Density | Final S. alterniflora Shoot Density | 14.2 | 1 | 0.0002 |
| Burrow Density | Final S. alterniflora Shoot Density | 0 | 1 | 0.953 |
| Year | Final S. alterniflora Shoot Density | 42.47 | 1 | 0 |
| Site | Final S. alterniflora Shoot Density | 1.31 | 1 | 0.2531 |
| Block | Final S. alterniflora Shoot Density | 1.76 | 1 | 0.184 |

**Table ST6:** Test statistics for a multivariate generalized linear model with a negative binomial error and log link comparing year, site, and block to initial living *Spartina alterniflora* live shoot density (shoots $0.125m^{-2}$) and year, initial crab density (crabs $0.25^{-2}$), initial sediment strength (kPa), initial living *S. alterniflora* live shoot density (shoots $0.125m^{-2}$), burrow density (burrows $0.25m^{-2}$), year, site, and block to final living *S. alterniflora* live shoot density (shoots $0.125m^{-2}$). These two models were evaluated for significance using Likelihood-Ratio (LR) chi-squared, Type II sums of squares.

**Supplemental Table ST7: Spartina alterniflora Biomass Model Test Statistics**

| Predictor | Response | LR Chisq | Df | p-Value |
|---|---|---|---|---|
| Initial Shoot Density | S. alterniflora Biomass | 0 | 1 | 0.9996 |
| Final Shoot Density | S. alterniflora Biomass | 1.48 | 1 | 0.2231 |
| Burrow Density | S. alterniflora Biomass | 0.01 | 1 | 0.9215 |
| Initial Sediment Strength | S. alterniflora Biomass | 20.76 | 1 | 0 |
| Initial Crab Dens. | S. alterniflora Biomass | 1.23 | 1 | 0.2665 |
| Final Sediment Strength | S. alterniflora Biomass | 0.5 | 1 | 0.4774 |
| Year | S. alterniflora Biomass | 1.35 | 1 | 0.2448 |
| Site | S. alterniflora Biomass | 6.96 | 1 | 0.0083 |
| Block | S. alterniflora Biomass | 38.72 | 1 | 0 |
| Initial-Final S. alterniflora Shoot Density Interaction | S. alterniflora Biomass | 21.76 | 1 | 0 |
| Final S. alterniflora Shoot Dens- Final Sediment Strength Interact. | S. alterniflora Biomass | 6.83 | 1 | 0.009 |
| Initial Crab Dens. Site Interaction | S. alterniflora Biomass | 5.36 | 1 | 0.0206 |

**Supplemental Table ST7:** Test statistics for a multivariate generalized linear model with a gamma error and log link comparing initial living *Spartina alterniflora* live shoot density (shoots 0.125m$^{-2}$), final living *S. alterniflora* live shoot density (shoots 0.125m$^{-2}$), fiddler crab burrow density (burrows 0.25m$^{-2}$), initial sediment strength (kPa), final sediment strength (kPa), year, site, block, a 2-way interaction term between initial and final living *S. alterniflora* live shoot density (shoots 0.125m$^{-2}$), a 2-way interaction term between final living *S. alterniflora* live shoot density (shoots 0.125m$^{-2}$) and final sediment strength (kPa), and a 2-way interaction term between initial fiddler crab density (crabs 0.25m$^{-2}$) and site to *S. alterniflora* aboveground biomass (g 0.125m$^{-2}$) measured at the end of the experiment. This model was evaluated for significance using Likelihood-Ratio (LR) chi-squared, Type II sums of squares.

**Supplemental Figure SF5: Initial and Final Shoot Density and S. alterniflora Comparison**

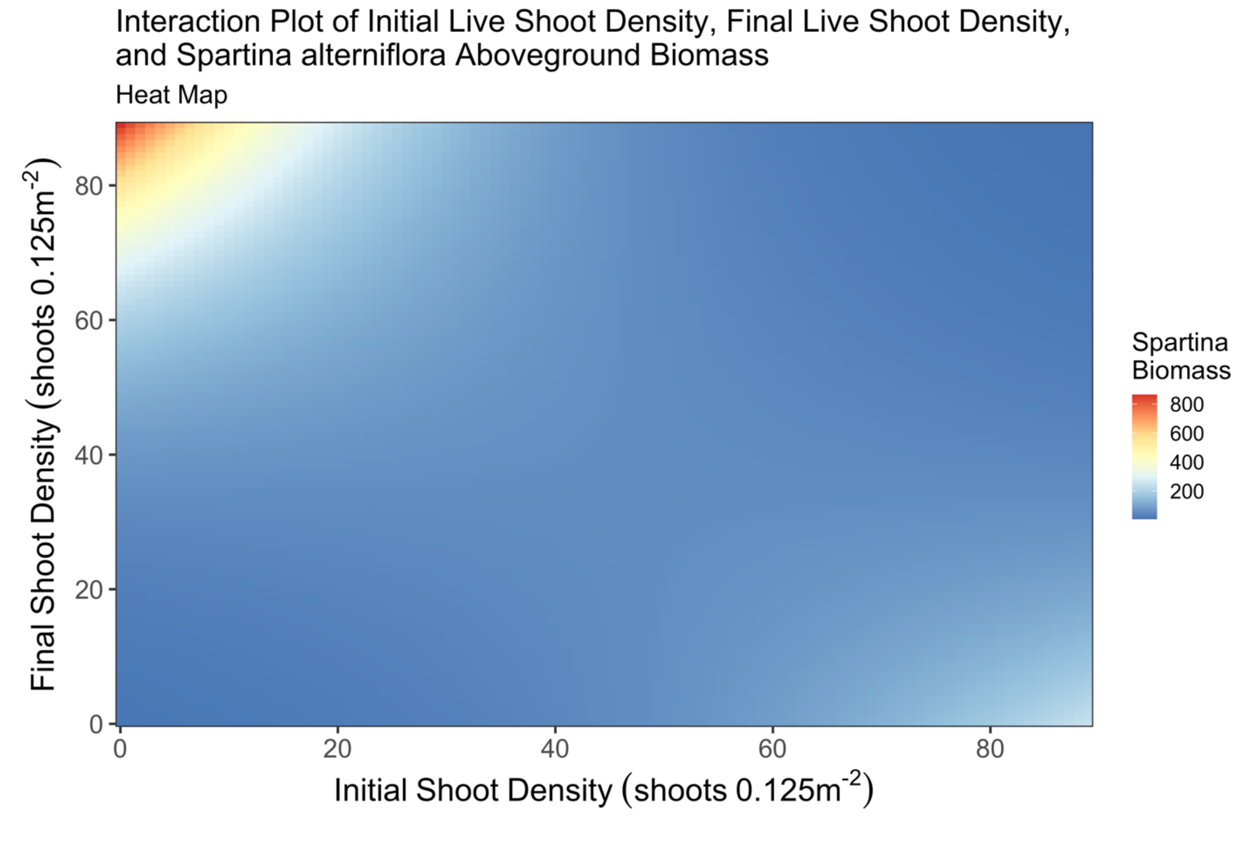


**Figure SF5:** Model generated *visreg* heat map showing how the interaction between initial and final *Spartina alterniflora* live shoot density (shoots $0.125m^{-2}$) causes changes in *S. alterniflora* aboveground biomass (g $0.125m^{-2}$). Warmer colors indicate higher biomass, cooler colors indicate lower biomass. The highest biomass was observed when initial living *S. alterniflora* live shoot density was low, and final living *S. alterniflora* live shoot density was high. This plot was drawn from the multivariate generalized linear model with a gamma error and log link comparing predictor variables and *S. alterniflora* aboveground biomass measured at the end of the experiment. For a complete list of predictors and test statistics, please see supplemental table ST7 above.

**Supplemental Table ST8: Decomposition Model Test Statistics**

| Predictor | Response | Sums Sq | Df | F Value | p-Value |
|---|---|---|---|---|---|
| Initial Sediment Strength | Decomposition | 0.09 | 1 | 5.42 | 0.0227 |
| Depth | Decomposition | 0.45 | 1 | 26.79 | 0 |
| Burrow Density | Decomposition | 0 | 1 | 0.03 | 0.8723 |
| Site | Decomposition | 0 | 1 | 0.08 | 0.7846 |
| Block | Decomposition | 0.05 | 1 | 2.99 | 0.0879 |
| Depth-Burrow Density Interaction | Decomposition | 0.14 | 1 | 8.45 | 0.0048 |
| Site-Block Interaction | Decomposition | 0.07 | 1 | 4.06 | 0.0476 |
| Site-Burrow Density Interaction | Decomposition | 0.12 | 1 | 7.1 | 0.0094 |
| Depth-Site Interaction | Decomposition | 0.05 | 1 | 3.19 | 0.0782 |
| Depth-Site-Burrow Interaction | Decomposition | 0 | 1 | 0.27 | 0.6017 |

**Table ST8:** Test statistics for a multivariate linear model with Gaussian error and an identity link comparing initial sediment strength (kPa), depth each litter bag was buried below the surface (cm), fiddler crab burrow density (burrows $0.25m^{-2}$), site, block, a 2-way interaction term between depth (cm) and burrow density (burrows $0.25m^{-2}$), a 2-way interaction term between site and block, a 2-way interaction term between site and burrow density (burrows $0.25m^{-2}$), a 2-way interaction term between depth and site, and a 3-way interaction term between depth, site, and burrow density (burrows $0.25m^{-2}$) to decomposition (g $litterbag^{-1}$). This model was evaluated for significance using Type II sums of squares F-test.

223 **Supplemental Table ST9: Site Comparisons Among All Models**

| Response Variable | Site Interactions | p-Value |
|---|---|---|
| Initial Sediment Strength | Year - Site | 0.062 |
| Initial Sediment Strength | Block - Site | 0.2322 |
| Final Sediment Strength | Burrow Density - Site | 0.3634 |
| Final Sediment Strength | Year - Site | 0.0997 |
| Final Sediment Strength | Block - Site | 0.9262 |
| Final Sediment Strength | Initial Shoot Density - Site | 0.2248 |
| Decomposition | Initial Sediment Strength - Site | 0.8597 |
| Decomposition | Depth - Site | 0.1352 |
| Burrow Density | Fiddler Crab Density - Site | 0.2655 |
| Burrow Density | Initial Sediment Strength - Site | 0.2756 |
| Burrow Density | Year - Site | 0.9125 |
| Burrow Density | Block - Site | 0.2595 |
| Shoot Density | Fiddler Crab Density - Site | 0.5553 |
| Shoot Density | Initial Sediment Strength - Site | 0.8731 |
| Shoot Density | Initial Shoot Density - Site | 0.3517 |
| Shoot Density | Burrow Density - Site | 0.7242 |
| Shoot Density | Year - Site | 0.2775 |
| Shoot Density | Block - Site | 0.1995 |
| Spartina Biomass | Initial Shoot Density - Site | 0.0797 |
| Spartina Biomass | Final Shoot Density - Site | 0.5296 |
| Spartina Biomass | Burrow Density - Site | 0.6663 |
| Spartina Biomass | Fiddler Crab Density - Site | 0.4248 |
| Spartina Biomass | Year - Site | 0.4425 |
| Spartina Biomass | Block - Site | 0.2999 |

**Table ST9:** Test statistics for multivariate linear and generalized linear models between predictors of ecosystem functioning, including year, block, burrow density (burrows $0.25m^{-2}$), initial and final *S. alterniflora* live shoot density (shoots $0.125m^{-2}$), initial and final sediment strength (kPa), and initial crab density (crabs $0.25m^{-2}$) with site to each ecosystem function response variable, including initial sediment strength, decomposition (g dried litter bag material$^{-1}$), burrow density, final live *S. alterniflora* live shoot density, and *S. alterniflora* aboveground biomass. Note, sediment strength, burrow density, and final live shoot density, while response variables of ecosystem function, are also predictors of other ecosystem functions included throughout our multivariate linear and generalized linear models. There were three instances of site interacting with an additional covariate; they were included in the main body of the text under the results section.
